# Comparison of subjective peripheral sensation, F-waves, and somatosensory evoked potentials in response to a unilateral pinch task measured on the contractile and non-contractile sides

**DOI:** 10.1101/2021.12.02.470947

**Authors:** Terumasa Takahara, Hidetaka Yamaguchi, Kazutoshi Seki, Sho Onodera

## Abstract

Depression of sensory input during voluntary muscle contractions has been demonstrated using electrophysiological methods in both animals and humans. However, the association between electrophysiological responses of the sensory system and subjective peripheral sensation (SPS) during a voluntary muscle contraction remains unclear. Our aim in this study was to describe the changes in SPS, spinal α-motoneuron excitability (F-wave to M-wave amplitude), and somatosensory evoked potentials (SEPs) during a unilateral pinch-grip task. Outcome variables were measured on the side ipsilateral and contralateral to the muscle contraction, and at rest (control). Participants were 8 healthy men, 20.9±0.8 years of age. The isometric pinch-grip task was performed at 30% of the maximum voluntary isometric force measured for the right and left hand separately. The appearance rate of the F-wave during the task was significantly higher for the ipsilateral (right) hand than for the contralateral (left) hand and control condition. Although there was no difference in F-wave latency between hands and the control condition, the amplitude of the F-wave was significantly higher for the ipsilateral (right) hand than for the contralateral (left) hand and the control condition. There was no difference in the amplitude of the SEP at N20. However, the amplitude at P25 was significantly lower for the ipsilateral (right) hand than for the contralateral (left) hand and the control condition. The accuracy rate of detecting tactile stimulation, evaluated for 20 repetitions using a Semmes–Weinstein monofilament at the sensory threshold for each participant, was significantly lower during the pinch-grip task for both the ipsilateral (right) and contralateral (left) hand compared to the control condition. Overall, our findings show that SPS and neurophysiological parameters were not modulated in parallel during the task, with changes in subjective sensation preceding changes in electrophysiological indices during the motor task. Our findings provide basic information on sensory-motor coordination.

## Introduction

When peripheral nerves are electrically stimulated, the ascending afferent input is projected to the somatosensory cerebral cortex via the spinal cord and the resulting cortical somatosensory evoked potentials (SEPs) can be recorded. During voluntary muscle contraction, sensory information induced by electrostimulation of the nerves supplying the contracting muscle is inhibited and the amplitude of SEPs decreases [1–4]. This suppression of the sensory potential is known as “gating.” The amount of gating during voluntary movement is dependent on the difficulty of the movement [5] and is observed during both the pre-movement and movement phases [6,7]. Based on these findings, the main functional role of gating is to eliminate unnecessary sensory information during the execution of purposeful voluntary movements.

Studies using the H-reflex [8] and F-wave [9,10] have shown an increase in the excitability of spinal α-motoneurons innervating active muscles during voluntary muscle contraction of the upper and lower limbs. The excitability of spinal α-motoneurons has also been shown to increase with contractions of distal [11] and contralateral [9] muscles. SEP gating has also been observed in the primary sensory cortex on the side contralateral to active muscle contraction [12], although there is no consensus on this finding [13]. Moreover, although depression of sensory input during voluntary muscle contraction has been demonstrated using electrophysiological methods in both animal and human studies, the association between the electrophysiological response of the sensory system and subjective peripheral sensation (SPS) during an active muscle contraction remains unclear.

In a previous study, we reported a reduction in cutaneous sensation on the dorsal surface of the hand during an isometric pinch-grip task under submaximal conditions compared to a no-motion (rest) condition [10]. However, it is not clear whether this response occurred locally only in the hand on the side of the contraction or would also be observed on the non-contracting side. Therefore, our aim in this study was to evaluate the changes in SPS, spinal α-motoneuron excitability, and SEPs on the side ipsilateral and contralateral to an active contraction of a hand muscle, the right abductor pollicis brevis (APB).

## Materials and Methods

### Participants and statement of ethics

The study group included 8 healthy adult men (mean ± SD age, 20.9± 0.8 years; height, 170.0 ±4.5 cm; and weight, 66.3 ± 9.5 kg) with no history of neurophysiological diseases. Our study was conducted in accordance with the principles of the Declaration of Helsinki. Informed consent was obtained from all participants. The study was approved by the Ethical Review Board of Kibi International University (No. 20-41).

### Electrical stimulation protocol

A square wave pulse, 0.2 ms in duration, was applied to the median nerve in the area of the carpal tunnel of the right hand, using a surface electrostimulation apparatus (NM-420S, Nihon Kohden, Japan), to stimulate the APB. The stimulation electrodes were placed 20 mm apart, with the cathode proximal and anode distal. The stimulation electrodes were secured using a fixation band to regulate the pressure applied to the electrodes. The minimum intensity of electrostimulation to induce an M-wave in the APB was confirmed.

### F-wave recording and analysis

Surface electromyogram (EMG) during electrostimulation was recorded over the right APB (side of stimulation at the carpal tunnel) using Ag/AgCl bipolar electrodes (5 mm diameter, 20 mm interelectrode distance; Nihon Kohden, Japan). The recording electrode was applied over the muscle belly of the APB, with the reference electrode placed over the first proximal phalanx. The position of both electrodes was fixed with surgical tape. Standard skin preparation was used prior for electrode placement: the site was cleaned with rubbing alcohol and the skin abraded using sandpaper to achieve a skin resistance of <5 kΩ. An EMG/evoked potential testing device (Neuropack MEB-9404, Nihon Kohden, Japan) was used. F-waves were recorded using a band-pass filter (1.5−3 kHz), at a sampling frequency of 10 kHz. To obtain the maximum M-wave, the intensity of electrostimulation was set to 1.2x the amplitude recorded for the M-wave appearance in the right APB. The stimulation frequency was set to 1 Hz and was applied for approximately 30 s. The following parameters of the F-wave were calculated during electrostimulation: appearance rate (%), latency (ms), and amplitude of the F/M ratio (%). The appearance rate was the number of F-waves observed on the monitor (threshold, 500 μV/D) from the total of the 30 possible waves generated by the electrostimulation. Latency was quantified as the average time from electrostimulation to F-wave initiation. The amplitude of the F-wave was expressed as the ratio of the average peak-to-peak amplitude of the F-wave to the maximum M-wave amplitude.

### SEP recording and analysis

Based on the international 10–20 system, the SEPs were recorded from the somatosensory area of the right upper arm (C3’, 2 cm posterior to C3) on the side ipsilateral to the electrostimulation. The reference electrode was placed at the point Fz. Electrodes were attached to the skin surface using a conductive paste, with a skin resistance of <5 kΩ after standard preparation. The electrostimulation intensity was set to just above the motor threshold, with a stimulation rate of 3 Hz. An EMG/evoked potential testing device (Neuropack MEB-9404, Nihon Kohden, Japan) was used and SEP waveforms were recorded using a band-pass filter (20 Hz to 10 kHz), at a sampling frequency of 10 kHz, with 200 responses averaged. SEP waveforms were evaluated for 100 ms, both at the time of electrostimulation and 100 ms after stimulation. Epochs with artifacts due to eye movement or blinking (> ±6 μV from baseline) were excluded automatically prior to averaging. A plate electrode was used to record the evoked electroencephalogram (Ag/AgCl electrode, NE-132B (Φ, 10 mm), Nihon Kohden, Japan). The peak-to-peak amplitude of the SPE at N20 and N20-P25, from baseline, which are early components after electrostimulation, were analyzed.

### SPS measurement

Prior to the experiment, the SPS threshold on the dorsal surface of the right hand was measured using the Semmes-Weinstein monofilament test (SAKAI Medical Co., Ltd., Tokyo, Japan) [10]. After establishing their peripheral sensory thresholds, participants reported the presence or absence of peripheral cutaneous stimulation during SPS measurements. Monofilament stimulation was performed using gradually thicker filaments, starting from thin filaments of 0.008 g. Confirmation tests were repeated approximately five times for each intensity and the filament thickness that could be sensed correctly at a rate of approximately 100% was defined as the SPS threshold. The experimenter lowered the filament vertically onto the hand, removed it, and returned it to its original position within 1 s. The stimulation interval of the filament was random and was repeated 20 times. Participants were instructed to give verbal cues when they sensed filament stimulation and the accuracy rate of detection was calculated. All measurements were performed by the same experimenter. The filament stimulation site was marked to avoid experimenter error resulting in random deviation in measurement due to a large stimulation site.

### Experimental procedure

Participants were seated in a chair with both arms placed on armrests, with eyes open to perform a pinch-grip task. When the pinch-grip task was performed with the right hand, the left arm and hand were maintained in a neutral position (no-motion, rest, condition). When the pinch-grip task was performed using the left hand, the right arm and hand were positioned in the neutral position. Prior to task performance, the maximum voluntary isometric force (MVIF) was measured separately for the right and left hand, from which the target force level was calculated. To calculate the MVIF, participants held the pinch force meter with the thumb and index finger and were asked to exert their maximum pinch force and to hold this force for 5 s. The peak force measured over this 5 s epoch was defined as the MVIF. After a sufficient rest period (≥10 min), the experimental task was performed. Participants were asked to maintain a 30% MVIF for a duration of 2 min, with the target pinch force to be exerted displayed visually on a computer screen placed 1 m in front of participants. The task was performed with both the right hand (ipsilateral to the side of recording) and the left hand (contralateral to the side of recording). In the control condition, no pinch force was exerted. The sequence of conditions was randomly selected across participants.

### Statistics

All values are presented as the mean ± standard deviation. Differences in measured parameters between conditions were evaluated using a repeated-measures analysis of variance. The sphericity of the data was evaluated using Mauchly’s test, with Greenhouse-Geisser-corrected significance values being used when sphericity was not met. Post-hoc analysis was performed using Tukey’s test for multiple comparisons. The statistical significance level was set at 5% (P<0.05) for all analyses. All analyses were performed using GraphPad Prism (version 8.3.1 for Machintosh).

## Results

A typical example of the M- and F-waves is shown in Fig 1A. The appearance rate of the F-wave (Fig 1B) was significantly higher for the side ipsilateral to the SEP recording (right hand, 85.1±11.3%) than for the contralateral side (left hand, 36.4±19.8%) and control (30.5 ±14.3%) condition (F_1.811,12.68_=39.78, P<0.01). There were no significant differences in the F-wave latency (Fig 1C) between the control condition (27.0±1.8 ms), the ipsilateral side (right hand, 26.3±1.5 ms), and contralateral side (left hand, 25.7±3.1 ms; F_1.523,10.66_=1.049, P=0.36). Similar to the appearance rate, the F-wave amplitude (Fig 1D) was significantly higher for the ipsilateral side (right hand, 7.4±4.4%) than for the contralateral side (left hand, 3.2±1.4%) and control condition (2.7±1.0%; F_1.060,7.419_=8.206, P=0.02).

**Fig 1.**
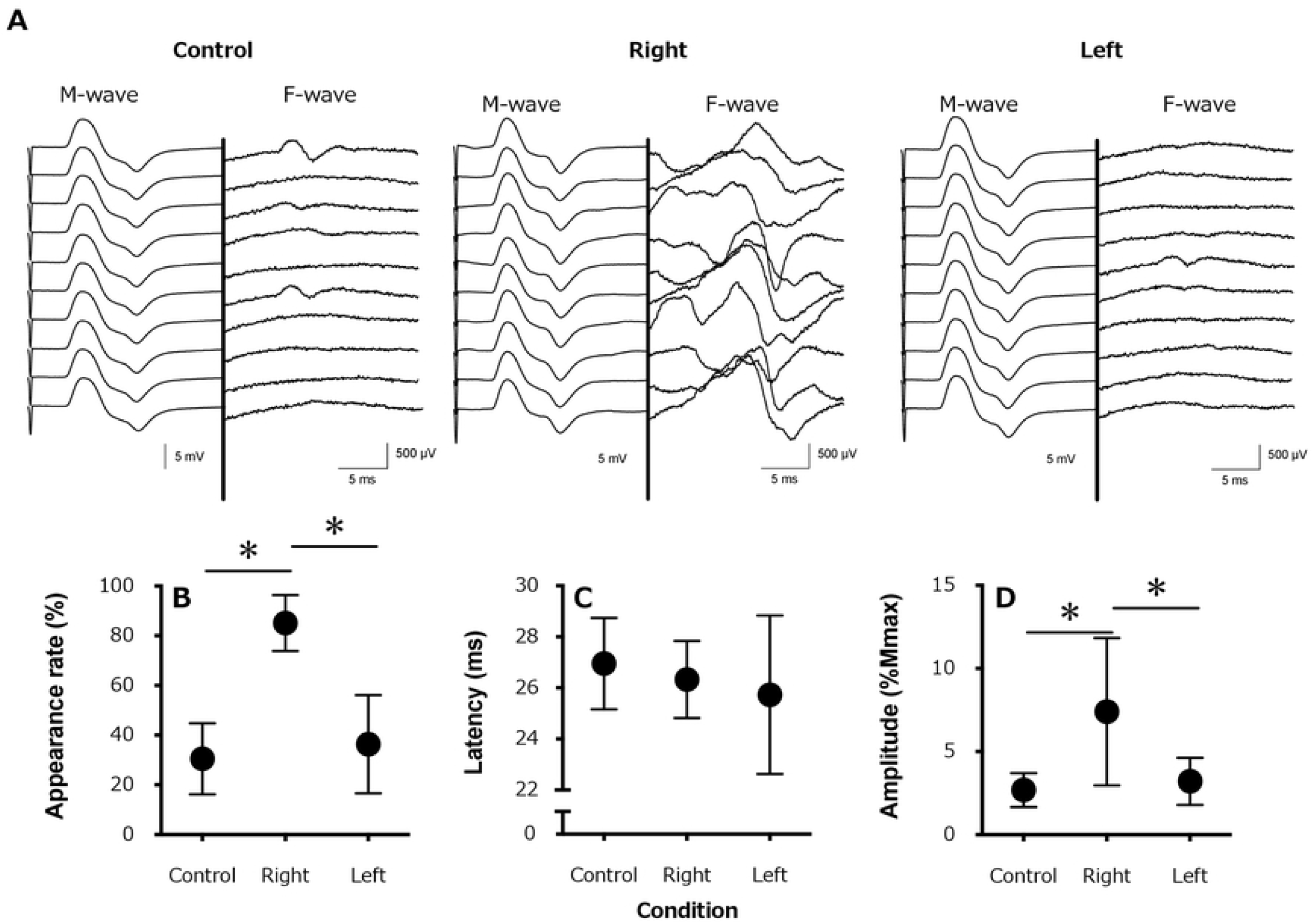
A typical example of an induced electromyogram waveform (10 stimulations) from a single participant in each condition. (**A**) The waveforms for the M-waves and F-waves are shown on the left and right side of the central thick line, respectively, with the amplitude scaling being 10 X higher for the F-than M-wave, for the control (rest), active contraction (right) side, and contralateral (left) side; (**B**) average appearance rate of the F-wave appearance; (**C**) F-wave latency, and (**D**) F-wave amplitude. *, P < 0.05

A typical example of SEP waveforms is shown in Fig 2A. There were no differences in the SEP amplitude at N20 (Fig 2B) between the control condition (3.41±1.11 μV), ipsilateral side (right hand, 2.73±1.16 μV) and contralateral side (left hand, 3.33±0.87 μV; F_1.247,8.726_=3.222, P=0.10). However, the SEP amplitude at P25 (Fig 2C) was significantly lower for the ipsilateral side (right hand, 4.65±1.27 μV) than for the contralateral side (left hand, 6.36±1.52 μV) and the control condition (6.42±1.22 μV; F_1.478,10.34_=14.63, P=0.001).

**Fig 2.**
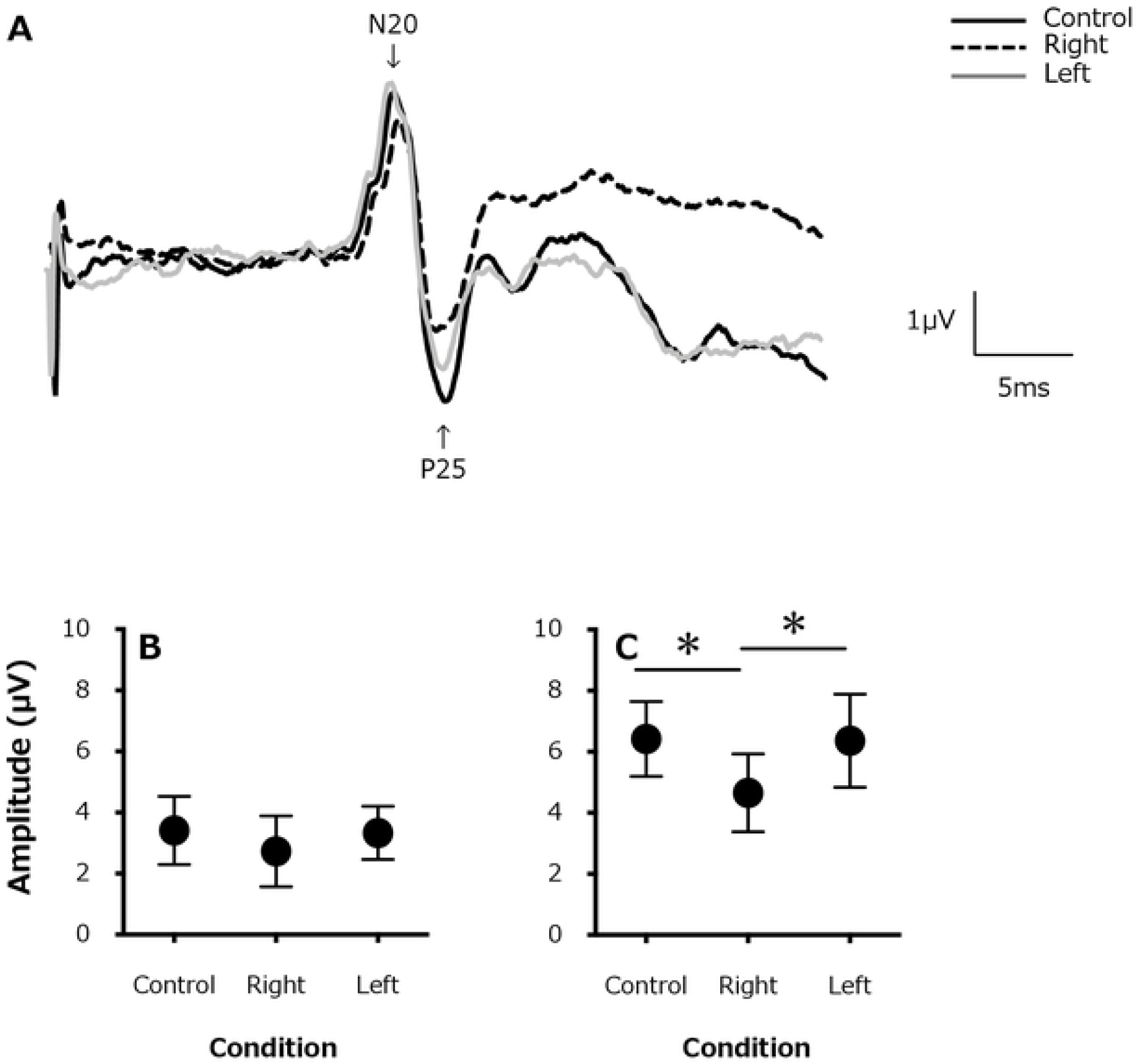
(**A**). A typical example of somatosensory evoked potentials (SEPs) waveforms from a single participant (for 200 stimulations) for the control (rest) condition (solid black line), in the active contraction (right) side (black dashed line), and contralateral (left) side (gray solid line). Changes in the average SEP amplitude (**B**) at N20 and (**C**) P25. *, P < 0.05

The accuracy rate for the 20 repetitions of monofilament stimulations was significantly lower for the ipsilateral (right) hand (61.9±21.9%) and the contralateral (left) hand (61.9±11.3%) than for the control condition (84.4±11.2%;F_1.297,9.080_=8.158, P=0.01; Fig 3).

**Fig 3.**
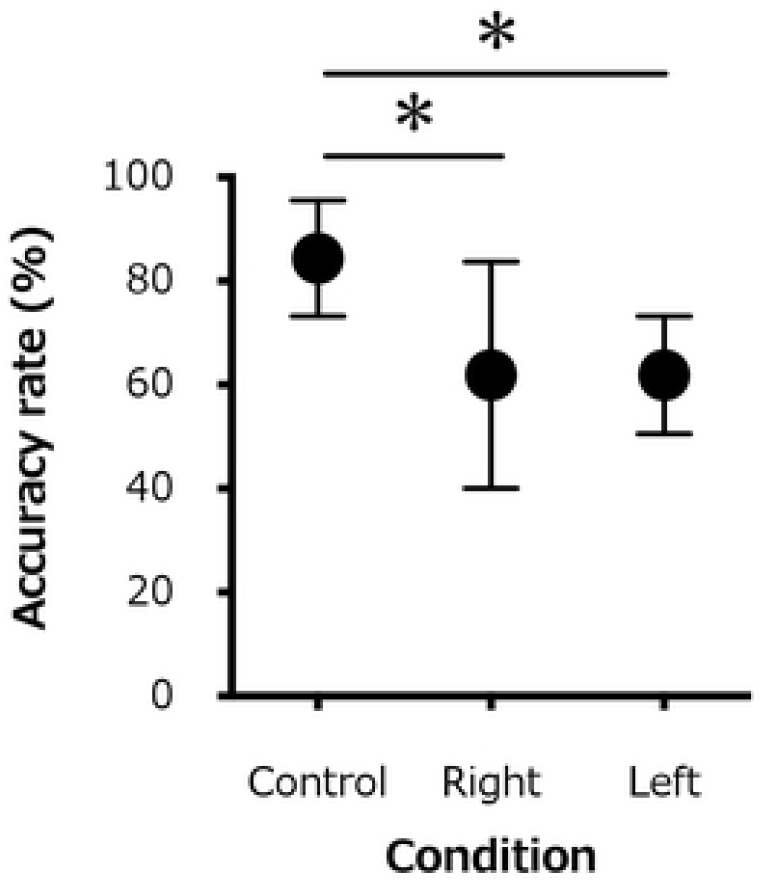
Changes in the average accuracy rate of detection for the 20 repetition monofilament test of all participants for the control (rest) condition, the active contraction (right) side, and the contralateral (left) side. *, P < 0.05

## Discussion

A novel observation of our study was that the pinch-grip task significantly reduced the SPS with active contraction of the right hand in both the ipsilateral (right) and contralateral (left) side compared to the control condition, although a significant increase in the appearance rate and amplitude of the F-wave and a significant decrease in the amplitude of the SEP (P25) were observed only on the ipsilateral side.

F-waves are muscle action potentials recorded when electrostimulation to a peripheral nerve causes retrograde conduction within the axon of an α-motoneuron, followed by subsequent anterograde conduction through the automatic firing of the α-motoneuron in the anterior horn of the spinal cord. In our study, the F-wave appearance rate for the right APB increased during the isometric pinch-grip task performed by the right hand (ipsilateral to the cortical recording side). The F-wave appearance rate indicates the number of motor units participating in the waveform [14] and naturally varies, even at rest. This natural variation suggests that sensory input influences the recruitment of α-motoneurons. This variation in the F-wave appearance rate increased during the isometric pinch-grip task in the ipsilateral (right) hand, which might reflect a suppression of activity within the corticospinal tract, which converges on the spinal anterior motor nerve, and inhibition via higher (cortical) control systems. This regulatory effect was not observed for contractions performed using the contralateral (left) hand, with no increase in the appearance rate of the F-wave for the left hand. This finding is different from previous reports of similar modulation of the F-wave on both the ipsilateral and contralateral side, with this difference likely reflecting differences in the motor task performed. While we used a pinch-grip task, previous studies used a hand-grip task, with the greater force generated by the hand-grip than the pinch-grip task increasing the firing rate and recruitment of α-motoneurons [15–17]. We do note that another study reported an increased responsiveness of neurons in the primary motor cortex for a precision (pinch-grip) rather than gross (power grip) motor task [18]. The increased muscle recruitment during a power grip, compared to a pinch-grip, task complicates the information measured from the upper motor centers due to the integrated processes of the central nervous system. These integrated processes exert an inhibitory effect on the α-motoneurons of the spinal anterior horn for muscles on the contralateral (non-contraction) side.

The source of the SEP at N20 is considered to be the 3b area of the primary somatosensory cortex, representing the stage when the sensory stimulation reaches the primary sensory cortex via the thalamus [19]. The source of the SEP at P25 is considered to be higher than the 3b area [20]. Therefore, the submaximal isometric pinch-grip task performed with the right hand in our study caused suppression of the ipsilateral somatosensory input at a higher level than the 3b area. Previous studies on SEP gating during voluntary movement have reported an absence of gating in components corresponding to N20, which is consistent with findings from previous studies. For these reasons, although the electrophysiological input that is projected to the primary somatosensory area is the same for a given amount of physical stimulation (regardless of the presence or absence of the motor task), this electrophysiological input is suppressed during the subsequent more complex phase of information processing. Additionally, as the amplitude of all components of the SEPs was not different between the left (contralateral) hand and the control (rest) condition, it appears that the electrophysiological sensory input is projected from the periphery to the primary somatosensory cortex, without influence from the contralateral muscle contraction.

In our study, we used the accuracy rate for cutaneous stimulation at the sensory threshold in the right hand as an index of SPS. The accuracy rate for cutaneous stimulation was reduced by submaximal isometric muscle contraction in both the ipsilateral (right) and contralateral (left) hands, compared to the control (rest) condition. This decrease in accuracy rate resulted from an increased difficulty in recognizing the sensory stimulation. There would be a need to develop a test in which the sensory information can be correctly recognized during a voluntary muscle contraction. Compared to the resting state, transient exercise has been shown to reduce skin temperature sensation [21], with the kinetic threshold also being lower during active muscle contraction than at rest [22]. Therefore, the tactile threshold of the skin surface may increase in the presence of a motor output, such as voluntary muscle contraction. Furthermore, the sensory thresholds with a muscle contraction may vary to a certain degree, both increasing or decreasing in amplitude. This phenomenon is also observed during muscle contraction on the contralateral side of the filament stimulation. These results suggest that localized muscle contraction modulates the SPS even in areas that are not related to a muscle contraction or movement.

The primary limitation to generalization of our findings is that the F-wave, SEP, and SPS measurements were conducted in separate sessions and not measured simultaneously in real time. As such, identification of the electrophysiological parameters corresponding to correct and incorrect results on the SPS test was not possible. Real-time measurement of F-waves, SEP, and SPS in future research would clarify the association between sensory-motor processes and subjective sensory changes in future studies.

## Conclusion

Overall, our findings show that SPS and neurophysiological parameters were not modulated in parallel during the task, with changes in subjective sensation preceding changes in physiological indices during the motor task. Our findings provide basic information on sensory-motor coordination.

## Author Contributions

Conceptualization: Terumasa Takahara

Data Curation: Terumasa Takahara, Hidetaka Yamaguchi, Kazutoshi Seki

Formal Analysis: Terumasa Takahara

Funding Acquisition: Terumasa Takahara

Investigation: Terumasa Takahara, Hidetaka Yamaguchi, Kazutoshi Seki

Methodology: Terumasa Takahara, Hidetaka Yamaguchi, Kazutoshi Seki

Project Administration: Terumasa Takahara

Resources: Terumasa Takahara, Hidetaka Yamaguchi

Supervision: Sho Onodera

Validation: Terumasa Takahara, Kazutoshi Seki

Visualization: Terumasa Takahara

Writing – Original Draft Preparation: Terumasa Takahara

Writing – Review & Editing: Terumasa Takahara, Sho Onodera

## References

1. Cheron G, Borenstein S. Gating of the early components of the frontal and parietal somatosensory evoked potentials in different sensory-motor interference modalities. Electroencephalogr Clin Neurophysiol. 1991;80: 522–530. doi: 10.1016/0168-5597(91)90134-j, PubMed PMID: 1720728.

2. Kakigi R, Koyama S, Hoshiyama M, Watanabe S, Shimojo M, Kitamura Y. Gating of somatosensory evoked responses during active finger movements magnetoencephalographic studies. J Neurol Sci. 1995;128: 195–204. doi: 10.1016/0022-510x(94)00230-l, PubMed PMID: 7738596.

3. Nakata H, Inui K, Wasaka T, Nishihira Y, Kakigi R. Mechanisms of differences in gating effects on short-and long-latency somatosensory evoked potentials relating to movement. Brain Topogr. 2003;15: 211–222. doi: 10.1023/a:1023908707851, PubMed PMID: 12866825.

4. Rushton DN, Rothwell JC, Craggs MD. Gating of somatosensory evoked potentials during different kinds of movement in man. Brain. 1981;104: 465–491. doi: 10.1093/brain/104.3.465, PubMed PMID: 7272711.

5. Kirimoto H, Tamaki H, Suzuki M, Matsumoto T, Sugawara K, Kojima S, et al. Sensorimotor modulation differs with load type during constant finger force or position. PLoS ONE. 2014;9: e108058. doi: 10.1371/journal.pone.0108058, PubMed PMID. PubMed Central PMCID: PMC4169486.

6. Bocker KB, Forget R, Brunia CH. The modulation of somatosensory evoked potentials during the foreperiod of a forewarned reaction time task. Electroencephalogr Clin Neurophysiol. 1993;88: 105-117. DOI: 10.1016/0168-5597(93)90061-s, PubMed PMID: 7681751.

7. Nishihira Y, Araki H, Ishihara A. Suppression of cerebral evoked potentials preceding rapid reaction movement. J Sports Med Phys Fitness. 1990;30: 291–296. PubMed PMID: 2266761.

8. Burke D, Adams RW, Skuse NF. The effects of voluntary contraction on the H reflex of human limb muscles. Brain. 1989;112: 417–433. doi: 10.1093/brain/112.2.417, PubMed PMID: 2706438.

9. Takahara T, Yamaguchi H, Seki K, Murata M, Onodera S. Effect of circulatory system response to motor control in one-sided contractions. Eur J Appl Physiol. 2018;118: 1773–1780. doi: 10.1007/s00421-018-3907-y, PubMed PMID: 29869712.

10. Takahara T, Yamaguchi H, Seki K, Onodera S. Sensory gating and suppression of subjective peripheral sensations during voluntary muscle contraction. BMC Neurosci. 2020;21: 41. doi: 10.1186/s12868-020-00592-2, PubMed PMID. PubMed Central PMCID: PMC7528260.

11. Zehr EP, Stein RB. Interaction of the Jendrassik maneuver with segmental presynaptic inhibition. Exp Brain Res. 1999;124: 474–480. doi: 10.1007/s002210050643, PubMed PMID: 10090659.

12. Staines WR, Brooke JD, Cheng J, Misiaszek JE, MacKay WA. Movement-induced gain modulation of somatosensory potentials and soleus H-reflexes evoked from the leg. I. Kinaesthetic task demands. Exp Brain Res. 1997;115: 147–155. doi: 10.1007/pl00005674, PubMed PMID: 9224842.

13. Tinazzi M, Zanette G, La Porta F, Polo A, Volpato D, Fiaschi A, et al. Selective gating of lower limb cortical somatosensory evoked potentials (SEPs) during passive and active foot movements. Electroencephalogr Clin Neurophysiol. 1997;104: 312–321. doi: 10.1016/s0168-5597(97)00023-3, PubMed PMID: 9246068.

14. Rivner MH. The use of F-waves as a probe for motor cortex excitability. Clin Neurophysiol. 2008;119: 1215–1216. doi: 10.1016/j.clinph.2008.01.103, PubMed PMID: 18406201.

15. Belhaj-Saif A, Fourment A, Maton B. Adaptation of the precentral cortical command to elbow muscle fatigue. Exp Brain Res. 1996;111: 405–416. doi: 10.1007/BF00228729, PubMed PMID: 8911934.

16. Cheney PD, Fetz EE. Functional classes of primate corticomotoneuronal cells and their relation to active force. J Neurophysiol. 1980;44: 773–791. doi: 10.1152/jn.1980.44.4.773, PubMed PMID: 6253605.

17. Maier MA, Bennett KM, Hepp-Reymond MC, Lemon RN. Contribution of the monkey corticomotoneuronal system to the control of force in precision grip. J Neurophysiol. 1993;69: 772–785. doi: 10.1152/jn.1993.69.3.772, PubMed PMID: 8463818.

18. Muir RB, Lemon RN. Corticospinal neurons with a special role in precision grip. Brain Res. 1983;261: 312–316. doi: 10.1016/0006-8993(83)90635-2, PubMed PMID: 6831213.

19. Allison T, McCarthy G, Wood CC, Darcey TM, Spencer DD, Williamson PD. Human cortical potentials evoked by stimulation of the median nerve. I. Cytoarchitectonic areas generating short-latency activity. J Neurophysiol. 1989;62: 694–710. doi: 10.1152/jn.1989.62.3.694, PubMed PMID: 2769354.

20. Allison T, McCarthy G, Wood CC, Williamson PD, Spencer DD. Human cortical potentials evoked by stimulation of the median nerve. II. Cytoarchitectonic areas generating long-latency activity. J Neurophysiol. 1989;62: 711–722. doi: 10.1152/jn.1989.62.3.711, PubMed PMID:2769355.

21. Gerrett N, Ouzzahra Y, Redortier B, Voelcker T, Havenith G. Female thermal sensitivity to hot and cold during rest and exercise. Physiol Behav. 2015;152: 11–19. doi: 10.1016/j.physbeh.2015.08.032, PubMed PMID: 26343771.

22. Taylor JL, McCloskey DI. Detection of slow movements imposed at the elbow during active flexion in man. J Physiol. 1992;457: 503–513. doi: 10.1113/jphysiol.1992.sp019390, PubMed PMID. PubMed Central PMCID: PMC1175743.

